# An optimized rabies vaccine vehicle for orotopical administration to wild vampire bats

**DOI:** 10.1101/2025.06.03.657068

**Authors:** Cole Knuese, Elsa M. Cárdenas-Canales, Travis McDevitt-Galles, María Magdalena Ramírez Martínez, Daniel Limonta, Lisa E. Powers, Daniel P. Walsh, Daniel G. Streicker, Jorge E. Osorio, Mostafa Zamanian, Tonie E. Rocke

## Abstract

Rabies vaccination of vampire bats (*Desmodus rotundus*) has been proposed as a superior control method to culling but has yet to be implemented. Success of rabies vaccination depends on a topical vehicle that spreads through a bat colony via allogrooming while additionally preserving vaccine immunogenicity. This work describes the *in vitro* and *in vivo* optimization of a new orotopical gel for rabies vaccine delivery to vampire bats. Autonomous transferability of our carboxymethyl cellulose (CMC) gel vaccine delivery formulation was tested in a microchipped vampire bat colony in rural Jalisco, Mexico. Intra-colony gel uptake was traced using the fluorescent biomarker rhodamine B. Importantly, application of topical treatment of ∼20% of the bat colony resulted in estimated uptake by over 85% of the colony. The *in vitro* stability of our raccoon poxviral-vectored rabies vaccine candidate within CMC was measured at time points up to 3 months at 40 °C, 23 °C, and 4 °C. Extended storage at 4 °C and short exposure at higher temperatures of potential vampire bat environments preserved vaccine titers within CMC. Furthermore, physical properties of our CMC formulation were compared to the previously used glycerin jelly at 40 °C, 20 °C, and 0 °C using rheological tests. These tests indicate that CMC exhibits optimal properties for topical application to bats even at extreme temperatures possible during field vaccination. This study advances our rabies vaccination strategy for vampire bats by providing a topical vehicle suitable for field application that may additionally be employed for other significant bat diseases.

## 1. Introduction

In Latin America, the vampire bat (*Desmodus rotundus*) is the primary reservoir of the deadly rabies virus (Rhabdoviridae, Lyssavirus)^1^. First reported in the early 1900’s, vampire bat-transmitted rabies (VBR) poses significant health risk to humans^2,3^ and, especially, to livestock^4^. Each year, VBR causes an estimated economic burden of $50 million USD to the livestock industry^5^. Additionally, the suitable habitat range for vampire bats is expected to spread due to changing climates^6,7^, and annual reports of VBR are trending upwards in several countries^1,8^. Recent models predict future expansion of the bats into the southern United States^7^ posing risk of extensive damages in this region with high populations of livestock^9^. Across the US, an estimated 40,000 individuals annually receive rabies prophylaxis post-exposure to natively found bat species^10^, and 90% of fatal rabies cases acquired from 2000-2022 were of bat origin^11^.

Attempts to mitigate VBR rely on vaccination of livestock and proactive or reactive culling of bat colonies via application of “vampiricide” – a topical paste containing poisons that spread by the allogrooming behavior of the bats^12^. However, since their inception in the 1970’s, these methods have been insufficient in preventing the expansion of VBR. Reactive application of vampiricide has been shown to exacerbate rabies virus transmission by provoking dispersal of infected bats to new roosts^13,14^. Furthermore, the poison may transfer to multiple non-target species in the wild, causing unintended mortality^1,12^.

The ingestion of topically applied (orotopical) vaccines has been previously proposed as a superior method for bolstering current control strategies^15,16^. Mathematical simulations predict that transferrable vaccines that similarly leverage bat allogrooming behavior outperform culling practices in reducing the probability, size, and duration of rabies outbreaks^17^. In earlier work, members of our group developed and tested a recombinant raccoon poxvirus (Poxviridae: Orthopoxvirus) expressing a mosaic lyssavirus glycoprotein (RCN-MoG)^18,19^. This vaccine was shown to produce protective immunity intradermally in A/J mice (*Mus musculus*)^18^, and orotopically in big brown bats (*Eptesicus fuscus*)^18^. Notably, orotopical vaccination blocked shedding of rabies virus in vampire bats^19^. Our team investigated glycerin jelly (GJ) as a topical vehicle to deliver the RCN-MoG vaccine in these captive bat colonies^18,19^ and to assess topical vehicle uptake in the wild^17^. However, the highly temperature dependent physical properties of GJ preclude its practicality for use in the field throughout Latin America, where tropical and subtropical climates commonly exceed its melting point^20,21^. Additionally, crystallization of GJ was noted in a separate field trial in Jalisco, Mexico^22^.

In this work, we describe the optimization of a novel carboxymethyl cellulose (CMC) gel vehicle for orotopical administration of the RCN-MoG vaccine to vampire bats. We compared physical properties of CMC gel and GJ with rheological tests to emulate application at environmentally relevant temperatures. Vaccine preservation was assessed by tracking viral titers over three-months across a range of temperatures. Lastly, we applied CMC gel mixed with the fluorescent biomarker rhodamine B (RB) to a wild colony of microchipped vampire bats in rural Jalisco, Mexico to investigate gel transfer and uptake of CMC in a field setting.

## 2. Methods

### Characterization of Physical Properties

#### Gel Preparation

Precise weights of CMC powder (Sigma-Aldrich, cat. no. 419281; St. Louis, Missouri, USA) were added to ultrapure water (Modulab® High Flow, Evoqua, Pittsburgh, Pennsylvania, USA) to prepare bulk solutions of 10%, 12.5%, and 15% (w/v). CMC gels were dissolved at room temperature overnight.

GJ (Carolina Biological Supply Co, cat. no. 865495; Burlington, North Carolina, USA) was melted at 37 °C and combined with ultrapure water at a ratio of 55.5:44.5 as was previously described^17^. GJ was solidified and kept at room temperature overnight.

#### Rheological Measurements

For characterization of rheological properties, an ARES LS2 Advanced research grade rheometer (TA Instruments, New Castle, Delaware, USA) was fitted with 50 mm cone and plate geometry. An FS18-HD Refrigerated/Heating Circulator (Julabo, Allentown, Pennsylvania, USA) and Peltier Plate were used for temperature control. Two different rheological tests were performed at 0 °C, 20 °C, and 40 °C with at least 5 min allowed for temperature equilibration. Testing was performed the day after gel preparation.

The first test, a steady rate sweep test, was used to record viscosity over a shear ramp from 0.05 s^-1^ to 200 s^-1^. Low shear rates characterize the near-zero shear rate and mimic behavior at rest. The range of 20 s^-1^ to 200 s^-1^ has previously been described as the shear rate to investigate spreading properties^23,24^.

Secondly, a dynamic strain sweep test characterized the viscoelastic properties of CMC and GJ. The elastic (storage) modulus G’ and viscous (loss) modulus G” were measured at a frequency of 1 hz with an initial strain of 0.015% up to 1000%.

### RCN-MoG Vaccine Stability Assay

#### Cells and Viruses

Vero cells (ATCC CCL-81) were cultured in Dulbecco’s Modified Eagle Media (DMEM; Sigma-Aldrich, St. Louis, Missouri, USA) supplemented with 5% fetal bovine serum and 1x antibiotics-antimycotics and incubated at 37°C with 5% CO_2_. The previously described RCN-MoG vaccine^18^ was used in the viral stability assay.

#### Vaccine Gel Preparation

To prepare RCN-MoG vaccine in CMC gel, RCN-MoG virus was homogenized in sterile 10mM Tris hydrochloride (Tris-HCl) buffer (pH 9.0) at a final concentration of 10^8^ PFU/mL in 4 mL total volume. CMC powder was then added at a final concentration of 12.5% (0.5 g) and stirred with a sterile microspatula.

For GJ preparation, RCN-MoG virus was added to sterile 10mM Tris-HCl buffer (pH 9.0). Then melted glycerin jelly was added such that the ratio of Tris buffer to glycerin jelly was 55.5:44.5 and the final concentration of RCN-MoG was 10^8^ plaque-forming units (PFU)/mL in 4 mL total volume. After homogenizing, 100 μL aliquots of the GJ preparation were transferred with a needle-free syringe into sterile 1.5 mL Eppendorf tubes.

Both the GJ aliquots and CMC gel were stored at 4°C overnight to allow the CMC powder to dissolve completely. Next, the CMC gel was briefly mixed and likewise aliquoted (100 μL) into sterile 1.5 mL Eppendorf tubes with a needle-free syringe. Aliquots were then placed at their respective storage temperatures (4 °C, 23 °C, or 40 °C). At each timepoint (24 hr, 48 hr, 96 hr, 1 wk, 2 wk, 1 mo, 3 mo), two aliquots were moved from each temperature and stored at -80 °C until titration by viral plaque assay.

#### Titration of RCN-MoG Vaccine by Viral Plaque Assay

Viral plaque assays were performed with Vero cells in 24-well plates (TPP, Trasadingen, Switzerland). Gel samples were thawed at 4 °C and dissolved by adding 900 μL of 1x DMEM to each tube with 1 hr incubation at 37 °C. Samples were mixed by pipetting every 15 min to facilitate dissolution. Cells were inoculated with 100 μL of ten-fold dilutions of the dissolved gels. After 1 hr of infection at 37 °C with 5% CO_2_, the inoculum was removed and 1 mL of plaque assay media (1x DMEM, 1.5% CMC, 2% FBS, 1x antibiotics-antimycotics) was added. Plates were fixed with 10% formaldehyde after 3 days of infection and stained with 1% crystal violet and 20% ethanol. Log fold change in viral titer was standardized to the initial time point (T=0).

### Western Blot

Six well plates of Vero cells were infected with 100 μL of dissolved CMC gel. After 18 hours, when >80% CPE was present, the supernatant and cells were harvested and centrifuged at 3,000 g for 10 minutes. After decanting, the cell pellet was resuspended in 100 μL of 4X Laemmli Sample Buffer (Bio-Rad, Richmond, California, USA) containing 10% 2-mercaptoethanol (Sigma Aldrich, Germany). Samples were denatured at 95°C for 5 min. SDS-PAGE resolved samples were then transferred to a nitrocellulose membrane (Bio-Rad, Richmond, California, USA). The membrane blocked was at 23 °C in 5% milk in PBS-T (blocking buffer) for 2 hours. Rabies virus glycoprotein polyclonal antibody (Invitrogen, Waltham, Massachusetts, USA) was used at 1:1000 in blocking buffer with 5% rabbit serum for 90 minutes at 23 °C. After three washes with PBS-T, Goat anti-Rabbit IgG (H+L) Secondary Antibody, HRP (Invitrogen, Waltham, Massachusetts, USA) was used at 1:4000 in blocking buffer with 5% goat serum for 90 minutes at 23 °C. After a final three washes, the membrane was developed with Pierce 1-step Ultra TMB Blotting solution (Thermo Fisher Scientific, Waltham, Massachusetts, USA). Membranes were scanned with an Epson Scanner (EPSON Perfection 4490 Photo) using the Epson Scan Utility v3.24 software.

### Bat Field Study

#### Ethics Statement

The Institutional Animal Care and Use Committee (IACUC) at the University of Wisconsin – Madison, WI, USA approved all animal work (Protocol No. V006701-A02). Permit number SPARN/DGVS/05928/24 was obtained from Secretaria de Medio Ambiente y Recursos Naturales (SEMARNAT, Mexico City, Mexico) for capturing and collecting hair samples from vampire bats.

### Study Site

The field study was performed at a vampire bat colony roosting in a long-abandoned ranch-house ∼10 km outside the Casimiro Castillo township, Jalisco, Mexico (19.580572, -104.534957). This colony had been monitored since October 2023 with follow-up captures every six months. During these previous captures, subcutaneous radio frequency identification (RFID) microchips were implanted in all captured bats to track bat movement at the roost through detections by a solar powered electromagnetic antenna and RFID detection system (Biomark, Boise, ID).

On October 31 and November 3 and 7, 2024, we performed diurnal captures. Two personnel entering the roost captured bats using butterfly nets, and a mist net was installed at the lone exit of the roost to ensnare bats attempting to fly out of the roost during capture efforts. Captured bats were immediately removed from the nets and kept in cages until processing. On each day of capturing, three capture sessions of 10 min spaced 1.5 hr apart were performed. Each captured bat was sexed and aged, with age determined by assessing fusion of the phalangeal epiphyses^25^. For analysis, subadults were grouped as juveniles.

Twenty of the bats were euthanized during captures 2 and 3 by local authorities as part of the activities of the Mexican rabies campaign for rabies prevention in livestock species due to equine rabies cases reported nearby.

### Gel Application and Sample Collection

The day prior to the first bat capture, 0.225 g (final concentration 0.3% w/v) RB was dissolved in 75 mL water. Then, 9.375 g CMC was added and manually stirred for a final concentration of 12.5%. The gel dissolved overnight.

During bat capture 1 (October 31, 2024), 1 mL of the gel was topically applied to the dorsal fur of bats with a needle free syringe. These bats were also given 100 μL orally to act as gel consumption controls for hair sampling. This dose of RB was previously determined to clearly mark vampire bat hair samples. The bats were then released back to the colony. On captures 2 and 3 (November 3 and 7, 2024), hair samples were plucked with tweezers from captured bats and stored in envelopes protected from light for later visualization with a fluorescence microscope.

### RB Visualization in Bat Hair

Uptake of RB within the hair samples was qualitatively assessed with a Zeiss Imager A2 microscope (Oberkochen, Baden-Württemberg, Germany) with a Zeiss 424931 Microscope Fluorescence Filter Cube (excitation 546 nm / emission 590 nm).

Observations were performed and assessed separately by two individuals. Samples were only recorded as positive if fluorescence was noted by both observers.

### Bat Colony Size Estimation

The colony size was estimated during capture 1 by using the Lincoln-Petersen method^26^ with the microchipped bats detected in the colony the night prior as the “marked” population. Each captured bat was scanned with an RFID scanner (AVID, Norco, California, USA) for microchip presence. This estimation method was used to determine the number of bats to topically treat during capture 1.

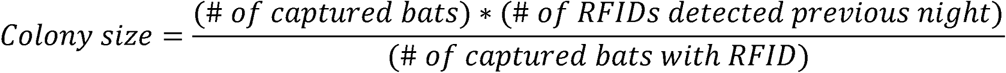

### Statistical Analysis

To assess variation in biomarker coverage across sex and age cohorts in our field trail we used a generalized linear model with a binomial distribution and a logit link function. For our response variable we used the presence of the biomarker (0/1) in the hair samples collected in unique bats from capture sessions 2 and 3. As our predictor variables, we included a grouping variable for both sex and age resulting in four distinct bat cohorts: adult female, adult male, juvenile female and juvenile male. The model was fit using the BRMS package^22^ in the R statistical language (R Core Team 2024). R version 4.4.2 was used to generate estimated biomarker coverage within the colony based on the positivity of collected hair samples. A hypothesis test built into the BRMS package was used to determine significant differences in the rates of coverage within the cohorts of non-treated bats.

For our covariate estimates, we used weakly informative priors with a normal distribution with a mean of 0 and standard deviation of 2. The model was run for 8,000 iterations with 2,000 iterations for warmup. To assess model convergence, we used the Gelman-Rubin test statistic, with a value <1.2 indicating convergence^27^ and visually inspected the trace plots for signs of divergent transitions.

Additionally, viral titer decay rates were modeled by linear regression and statistical comparisons were made using emtrends function of the emmeans R package^28^. A Chi-squared test was used to analyze the age and sex distribution of bats across the three days of captures. Fisher’s exact test was used to analyze differences in RB positivity of collected hair samples of bat cohorts stratified by the combination of age and sex.

## 3. Results

### CMC-based vaccine formulation exhibits optimal properties for topical application

To compare the consistency of CMC and GJ across a range of temperatures, rheological testing was performed. The temperatures tested (0 °C – 40 °C) represent extreme conditions under which a vaccine gel might be applied to vampire bats for rabies vaccination or to other bat species in temperate climates.

A steady rate sweep test was performed to measure viscosity across a range of shear rates, or rates of deformation. As the viscosity of CMC solutions is modulated by concentration (Figure 1A; Figure S1), a concentration of 12.5% was considered to have the optimal consistency. The apparent zero-shear viscosity of 12.5% CMC, estimated by the Newtonian plateau and an approximation of gel behavior at rest, exhibited much variation with changes in temperature compared to GJ (Figure 1B). Zero-shear viscosity for 12.5% CMC ranged from approximately 110-400 Pa*s from 0 to 40 °C compared to 0.05 – 1,070 Pa*s for GJ. At all temperatures, the viscosity of CMC and GJ decreased as shear rate increased, indicating shear-thinning behavior where viscosity decreases as deformation rates increase.

**Figure 1.**
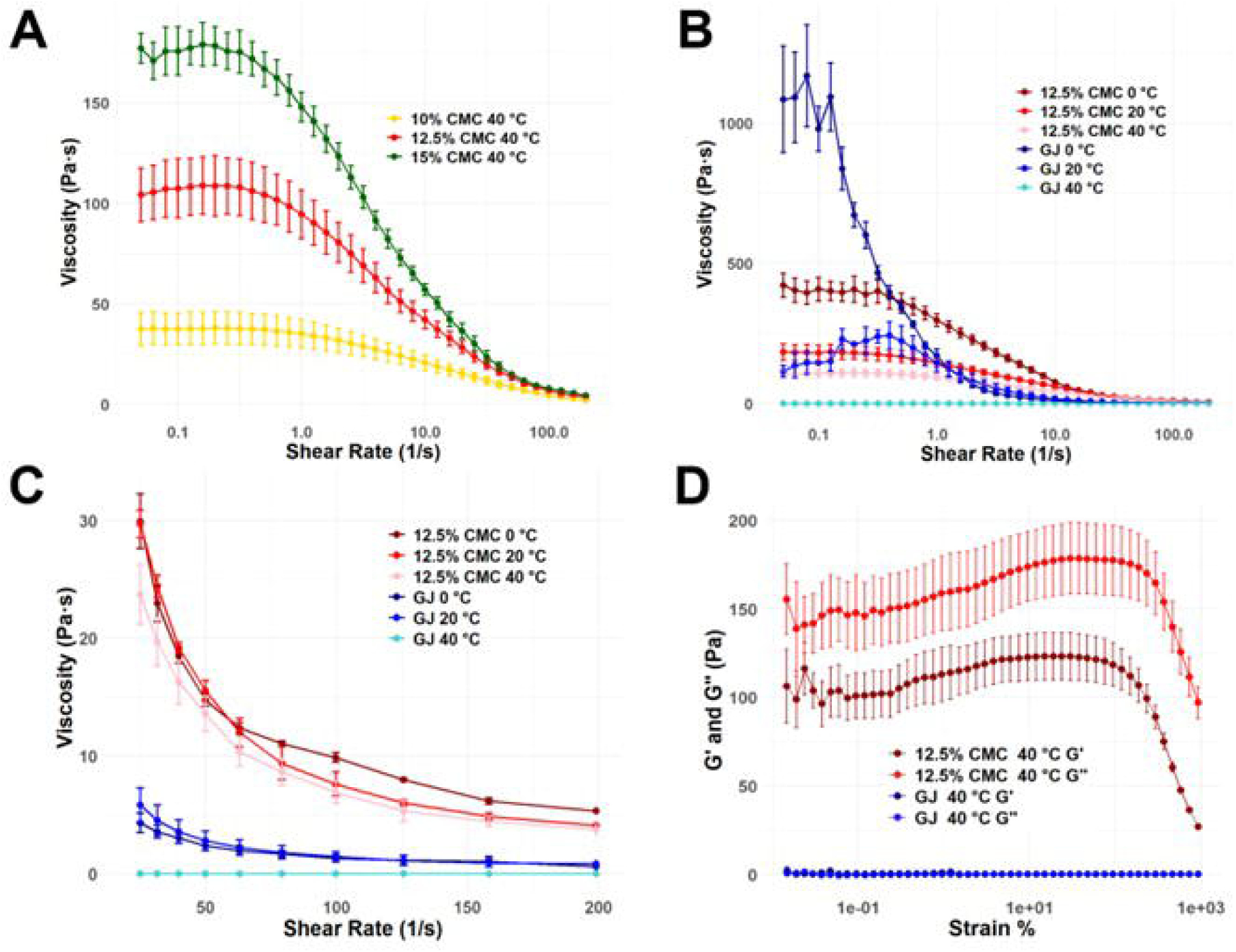
Rheological properties of carboxymethyl cellulose (CMC) gel and glycerin jelly (GJ). **A)** The viscosity of 10%, 12.5% and 15% solutions of carboxymethyl cellulose (CMC) at 40 °C and **B)** the viscosity of 12.5 % CMC and GJ at 0 °C, 20 °C, and 40 °C were measured by a steady rate sweep test. Panel **C)** displays the viscosity of CMC and GJ depicted over shear rates associated with spreading (20 – 200 s^-1^). **D)** Viscoelastic properties of CMC and GJ at 40 °C were measured by a dynamic strain sweep test. G’ represents the elastic modulus – energy stored during deformation, while G” represents the loss modulus – energy dissipated during deformation. Error bars represent standard error of three experimental replicates.

At shear rates within the range of 20-200 s^-1^ (Figure 1C), shear rates associated with spreading^23,24^, CMC was more viscous than GJ and varied minimally with changes in temperature. In the dynamic strain sweep test, CMC gel exhibited viscoelastic properties and acted as a viscoelastic liquid even at 40 °C (Figure 1D). GJ which displayed properties of an elastic solid at 0 °C and 20 °C (Figure S2) but became liquified at 40 °C (Figure 1D).

### CMC gel preserves viral vaccine titers

We measured viral titers at time intervals up to three months to assess preservation of our RCN-MoG vaccine within CMC and GJ under different temperature conditions. Stability of the RCN-MoG vaccine titers was highly temperature dependent within both CMC and GJ formulations (Figure 2A-C). High titers were preserved up to 3 months at 4 °C (Figure 2B) with minimal decay of RCN-MoG in both formulations (CMC: -0.0385 ± .0221 PFU/mL per week; GJ: -0.00311 ± 0.0191 PFU/mL per week; Figure 2C). At 23 °C, viral titer decreased in CMC at a moderate rate (-0.463 ± 0.062 Log10 PFU/mL per week), while in GJ decay was slower (-0.157 ± 0.041 Log10 PFU/mL per week). RCN-MoG viral titer rapidly decreased at 40 °C in both CMC (-3.50 ± 1.03 Log10 PFU/mL per week) and GJ (-1.50 ± 0.28 Log10 PFU/mL per week). RCN-MoG was no longer detectable at 40 °C after 30 days in CMC and after 90 days in GJ. Of note, slightly more virus was recovered from dissolved GJ (Log10 8.02 +/- 0.123 PFU/mL) than CMC (Log10 7.64 +/- 0.136 PFU/mL) at the initial time point (Figure S3).

**Figure 2.**
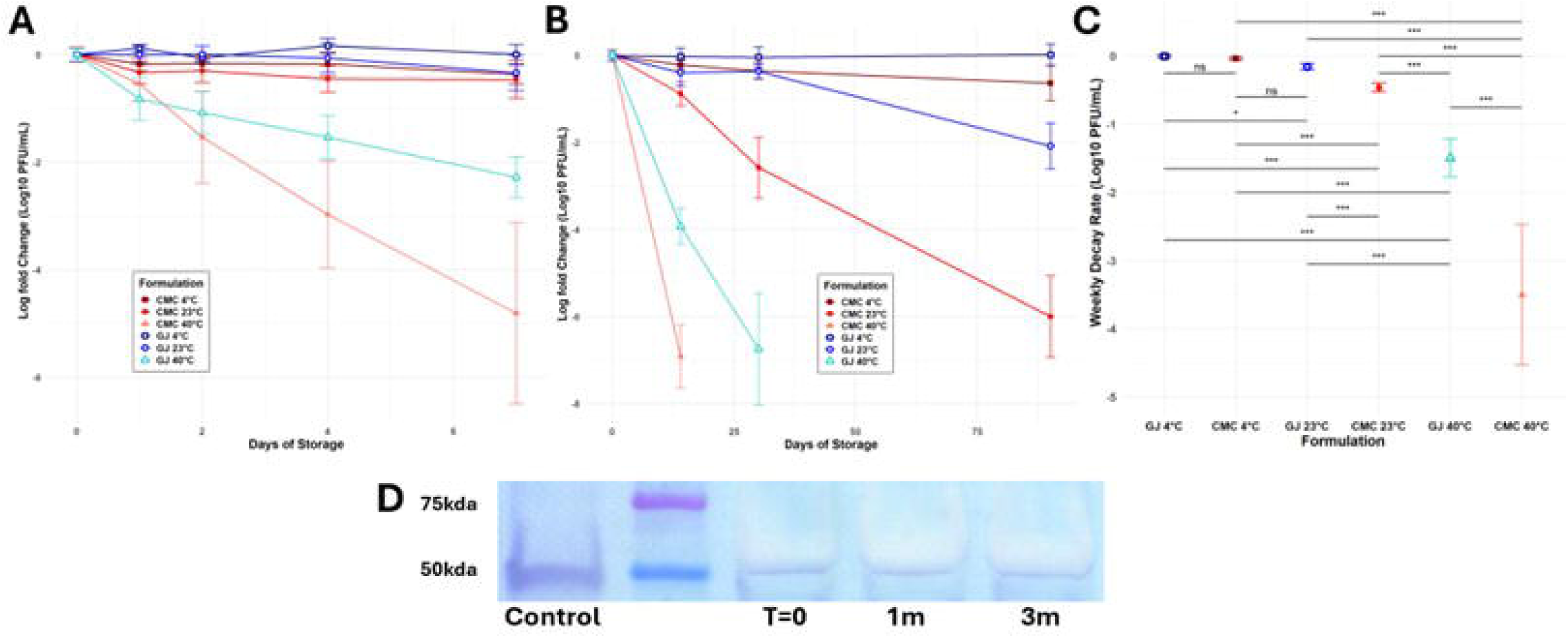
Viral stability of a recombinant raccoonpox vaccine expressing a mosaic lyssavirus glycoprotein (RCN-MoG) stored within carboxymethyl cellulose (CMC) and glycerin jelly (GJ) formulations at 4. °**C, 23** °**C, and 40** °**C.** Log fold change in vaccine titer measured over **A)** seven days and **B)** three months of storage by plaque assay in Vero CCL-81 cells at each time point after three days of infection. Error bars depict the standard error of three experimental replicates. **C)** Weekly decay rates of RCN-MoG titer within CMC and GJ at each storage condition were determined by a linear regression model. Statistical comparisons were made using emtrends (ns = no significance; * = p < 0.05; *** = p < 0.001). **D)** MoG antigen expression of cell lysates infected with RCN-MoG after storage in CMC at 4 °C for one and three months was detected by western blot. Cells were infected with 100 μL of dissolved gel and protein was collected 18 hours post-infection.

Cells infected with RCN-MoG stored in CMC at 4 °C for three months still produced antigen detected by western blot (Figure 2D). While replication levels of samples stored at 23 °C reduced drastically over time, vaccine poxviral DNA was preserved throughout the study (Table S1-S2; Figure S4).

### High uptake rate of CMC-based delivery formulation within a wild vampire bat colony

CMC gel uptake by wild vampire bats was measured using the fluorescent biomarker rhodamine B (RB) within a microchipped colony in Jalisco, Mexico (Figure 3A). The bats inhabited an abandoned ranch-house (Figure 3B) in which we previously installed an RFID system for microchip detection (Figure 3C). Over three capture sessions (Figure 3D), 156 total *Desmodus rotundus* bats were captured (Figure 3E), representing 95 unique bats. No bats of other species were captured.

During capture 1, 47/89 (52.8%) bats had RFID microchips from previous trips to the site (Figure 3E), while 62 microchipped bats were detected the night before capturing (Figure 3F). This results in an estimated colony size of 117.4 (± 5.625 SE) by the Lincoln-Petersen method. One milliliter of 12.5% CMC with 0.3% rhodamine B (RB) was topically applied to the dorsal fur of 24 bats (13 females and 11 males), representing an estimated 20.5% of the colony. Forty-two new microchips were implanted during capture 1, resulting in the observed increase in detected RFIDs (Figure 3F).

Age and sex demographics within the entire colony were determined from the proportions of unique bats captured belonging to each subpopulation across the three captures. Of the bats in the colony, 57/95 were female (60.00%), and 66/95 were adults (69.47%) (Table 1). RFID data shows a decline in the number of microchipped bats detected each night within the colony beginning after capture 2 (Figure 3F). The age and sex distribution of bats captured during captures 2 and 3 were not statistically different from capture 1 (Chi-squared test, p = 0.165).

**Table 1.** Age and sex distribution of unique captured vampire bats (*Desmodus rotundus*) during three capture sessions in Jalisco, Mexico. Age was determined by fusion of the phalangeal epipyhyses. Subadults were grouped with juveniles.

| **Cohort** | **Unique Bats Captured** | **Percentage of Captures** |
| --- | --- | --- |
| Adult Female | 40 | 42.11% |
| Adult Male | 26 | 27.37% |
| Juvenile Female | 17 | 17.89% |
| Juvenile Male | 12 | 12.63% |
| Total | 95 | 100% |

**Figure 3.**
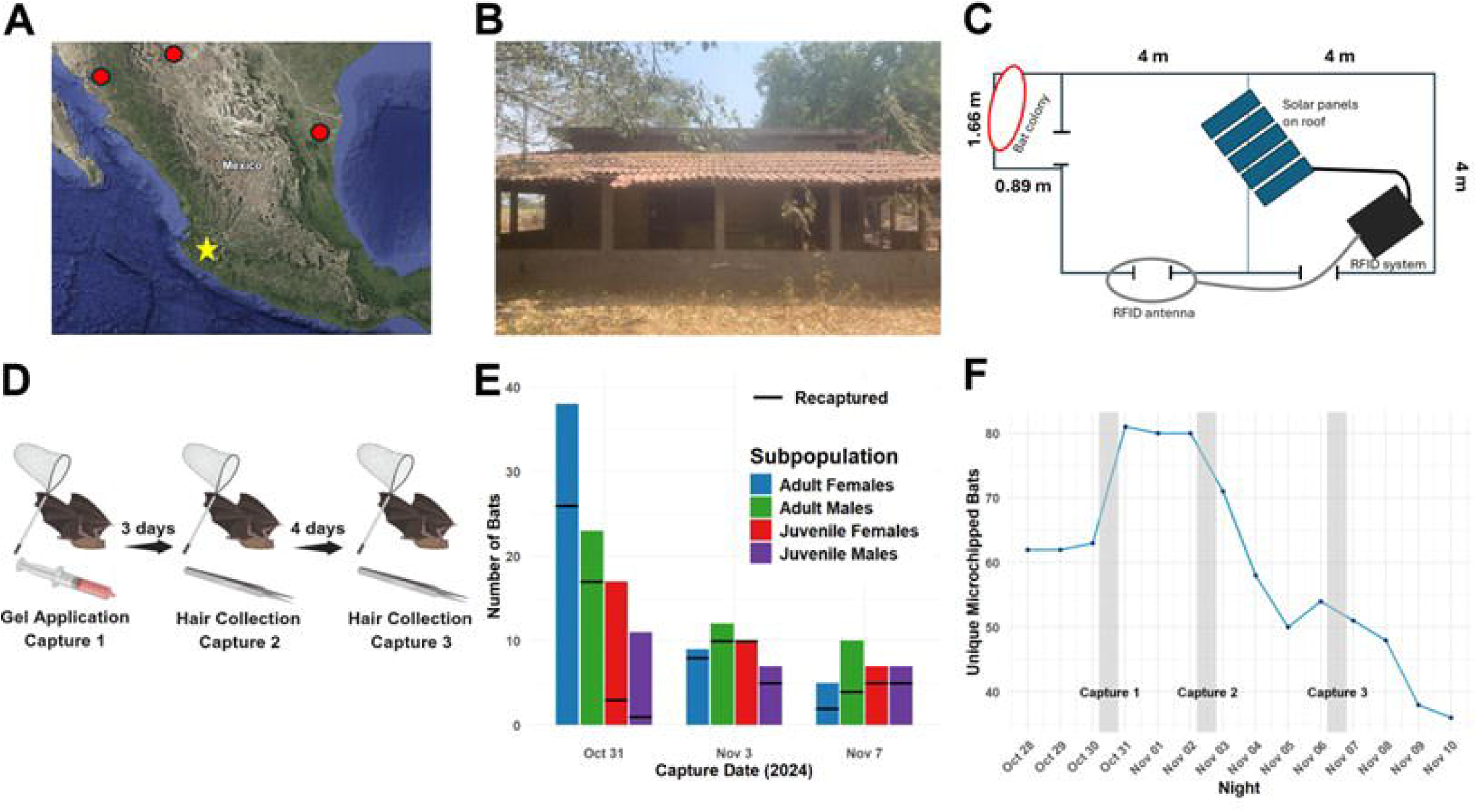
Study design and demographics of captured vampire bats (*Desmodus rotundus*) during three capture sessions in Jalisco, Mexico. A) In Mexico, the field study location is marked with the yellow star, while regions with the northernmost sightings of vampire bats are labeled with red circles (map retrieved from Google Earth v 10.80.0.1 on May 15, 2025). **B)** The vampire bat colony inhabited the photographed abandoned ranch-house. Photograph by C.K. on May 28, 2024. **C)** In the ranch-house, a radio frequency identification (RFID) system was installed (overhead schematic) to detect movement of microchipped bats. **D)** Three captures were performed for gel application with rhodamine B biomarker during capture 1 and hair collection during captures 2 and 3. **E)** During each capture, bats were sexed and aged. Lines represent the number of recaptures from the previous capture session, or in the case of capture 1, bats with RFID microchips from previous captures at this site. **F)** Each night, the RFID system detected the movement of microchipped bats into and out of the roost. The y-axis represents the number of unique RFIDs detected each night (x-axis).

During bat capture sessions 2 and 3, 34 and 27 hair samples were collected, respectively. Accounting for recaptures, hair samples were obtained from 48 unique bats, including 13 that were initially topically treated with CMC gel (Figure 4A-D). Hair samples were visualized with a fluorescent microscope (Figure 4E-F), and all 13 treated bats that were recaptured were RB-positive. Of the nontreated bats, 29/35 (82.86%) were RB positive. Notably, the only negative hair samples belonged to adult male bats resulting in a statistically different proportion of positive samples compared to the other cohorts (Fisher’s exact test, p = 0.002). Three adult male bats that were captured during both captures for hair sample collection had positive hair samples on capture 2 but negative samples on capture 3 (Figure 4A). These individuals were still considered RB positive for analysis.

Based on the proportion of positive and negative hair samples and colony demographic estimates, total gel uptake by the colony was modeled using Bayesian regression modelling with the BRMS package^29^. Gel uptake within non-treated and non-captured bats was predicted using the brm model (Figure 4G). Near 100% rates of uptake were predicted in non-treated adult female (97.4%; confidence interval (CI): 85.7% - 100%), juvenile female (94.5%; CI: 73.3% - 100%) and juvenile male (98.8%; CI: 92.3% - 100%) cohorts. Notably, an estimated uptake of 48.4% (CI: 30.4%-69.6%) was found in non-treated adult males. Across the entire colony, uptake was estimated at 88.0% (103/117; 90 - 110) (CI: 76.9% - 94.0%). Based on this projection, 103 bats consumed the gel, with each bat consuming on average ∼200 μL of the total 24 mL of topically applied gel (1 mL applied to 24 bats), assuming all the gel was consumed.

**Figure 4.**
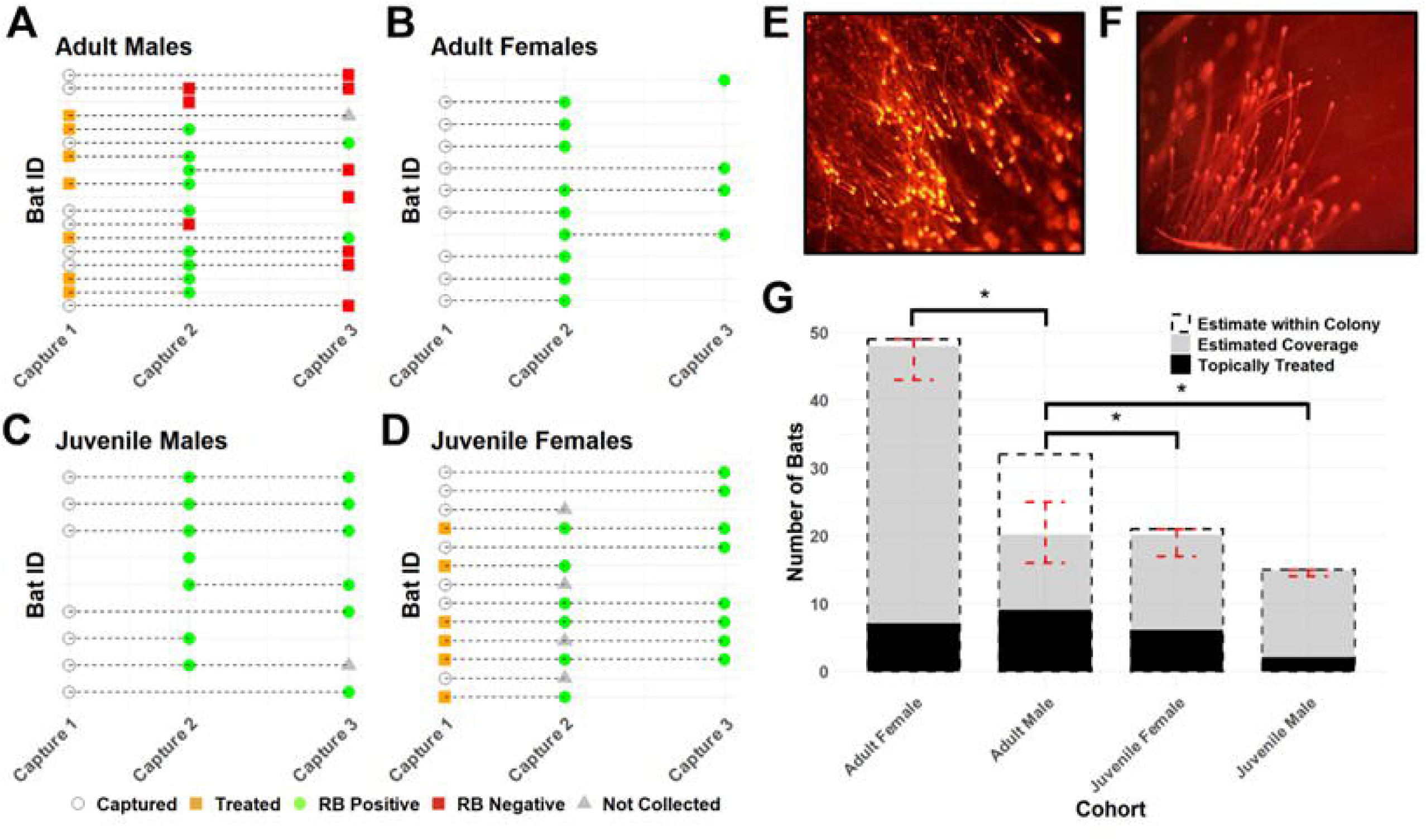
Estimated gel uptake within a vampire bat (*Desmodus rotundus*) colony in Jalisco, Mexico, after approximately 20.5% of the colony was treated topically with 1 mL of carboxymethyl cellulose (CMC) gel containing 0.3% rhodamine B (RB). Panels **A** (adult males), **B** (adult females), **C** (juvenile males), and **D** (juvenile females) depict capture and treatment status during capture 1 and collection and positivity of hair samples of bats captured during captures 2 and 3. During capture 1, orange squares represent topically treated bats, while open circles were captured, but not treated. The presence of orange fluorescence was used to distinguish between **E)** positive and **F)** negative hair samples. Hair samples were visualized with a Zeiss Imager A2 microscope (Oberkochen, Baden-Württemberg, Germany) and a Zeiss 424931 Microscope Fluorescence Filter Cube (excitation 546 nm / emission 590 nm). **G)** A Bayesian model was used to calculate estimates and confidence intervals of estimated coverage within each subpopulation of bats. Statistical significance indicates difference in coverage of non-treated bats and was determined by brms hypothesis testing.

## 4. Discussion

While rabies vaccination of vampire bats in captivity has been shown to reduce rabies virus shedding^19^ the delivery of the vaccine to wild vampire bats presents a major barrier to the implementation of this strategy as a rabies control measure. Although effective in humans and domesticated animals, parenteral vaccination is impractical for bats. Likewise, oral baits successfully used for other wildlife species^30^ are not suitable for bats due to differing feeding behavior. Because topically administered poisons demonstrate high rates of oral uptake within bat colonies^31,32^, orotopical vaccination of vampire bats has been proposed but lacks an optimal delivery vehicle. In this work, we describe the development of a topical vehicle that maintains optimal physical properties even at extreme temperatures, preserves our viral vector vaccine candidate, and exhibits high rates of transfer and oral uptake between treated and untreated bats in a wild bat colony.

Previous bat studies tested petroleum jelly^33^ – a paste commonly used for anticoagulant poison delivery – or glycerin jelly (GJ)^17–19^ as the topical vaccine delivery vehicles. Petroleum jelly proved difficult to work with resulting in dilution of the vaccine and substantial loss of vaccine paste^33^. Meanwhile, the consistency of GJ is highly temperature dependent as demonstrated by our study. We found the mixture of GJ in water previously used^17^ has a melting temperature around 30 °C. While vampire bats typically roost where temperatures remain below 37 °C^34^ and do not well tolerate temperatures above 33 °C^35^, environmental temperatures may exceed 40 °C during heat waves in some regions^20^.

Characterization of physical properties of a topical vehicle under environmental temperatures is necessary for proper application as revealed by the shortcomings of GJ. Rheology, measuring how a material sample responds to deformation, presents a useful method for measuring physical properties of gel-like materials^36^. The rheological properties of CMC solutions are well studied and vary with the molecular weight and degree of substitution of the CMC powder used^37–39^. The CMC gel preparation used in our study demonstrates shear thinning behavior and viscoelastic properties corresponding with these previous reports. In a steady rate sweep test, CMC demonstrates less variation than GJ in zero-shear viscosity over the studied range of temperature. At 40 °C, the near-zero shear rate of CMC was approximately 110 Pa*s, demonstrating high viscosity at high temperatures. This improves on the GJ formulation which has a near-zero shear rate approaching 0 Pa*s at 30 °C when it becomes completely liquified. In the characterization of viscoelastic materials, the storage modulus G’ characterizes the elastic component (energy stored during deformation) and the loss modulus G” characterizes the viscous component (energy dissipated during deformation). In CMC, G” is greater than G’ in the linear viscoelastic region which suggests CMC acts as a viscoelastic liquid that flows slowly over time. This may contribute to improved surface wettability and to the enhanced stickiness of CMC that we observed compared to GJ. In GJ, G’ > G”, indicating primarily elastic solid-like behavior. Notably at 0 °C, G’ becomes greater than G” in CMC indicating trends toward solidification at temperatures near freezing, while still exhibiting viscoelastic properties.

Additional described properties of CMC support its use as a topical vaccine vehicle. The muco-adhesion of CMC^40^ may enhance viral entry into mucosal tissue. Mucoadhesive cellulose-based films have been previously used to facilitate entry of Modified Vaccinia Ankara – a commonly used poxviral vaccine vector - into the buccal mucosa of mice^41^. CMC is also commonly used in drug delivery^42^, is non-toxic at high quantities^43^, and was found to mix easily with buffer containing RB.

To facilitate vaccination strategies, the vaccine must be readily shipped and stored without dependence on cold-chains in regions where reliability of infrastructure may be unpredictable^44^. Our assessment of viral stability in CMC under different temperature conditions provides insight into vaccine preservation under storage conditions and persistence in field-relevant conditions. Minimal decay of viral titer within CMC was evident at refrigeration temperatures (4 °C) with only a 0.63 log fold decrease in viral titer after 3 months of storage. This will enable shipment and storage of the vaccine when challenges in infrastructure prevent maintaining temperatures below freezing. If refrigeration is lost, vaccine titer may also be preserved up to 1 week at 23 °C with only a 0.46 log fold decrease in titer and 24 hrs at 40 °C with 0.54 log fold decrease.

Although virus stability was generally better preserved in GJ;, the viral titer initially decreased more rapidly in GJ than in CMC at 40 °C with a log fold decrease of 0.82 after 24 hours. Other additives could be considered to improve stability within the CMC gel formulation such as sucrose, which has been shown to enhance poxviral vaccine vector stability^45^. The implications of these vaccine preservation data depend on the determination of the minimum effective dose of RCN-MoG in vampire bats; however, during field application, we expect gel to be consumed within hours after application.

Like applications of vampiricide, orotopical vaccination utilizes the natural propensity of vampire bats to allogroom to facilitate the transfer of vaccine between treated and untreated bats. Ideally, only a small portion of a colony would need to be treated to effectively manage rabies due to difficulties in locating and treating entire roosts.

Vampire bat captures are commonly performed at ranches where bats are known to be feeding to apply vampiricide, rather than at the roost istelf^31^. This highlights the importance of understanding the rates of vaccine transfer and uptake within colonies and between cohorts at low rates of application. The RB biomarker used in this study is a common proxy for measuring uptake of treatment in wildlife species given its properties to stably mark the hair follicle and shaft with a fluorescent signal^17,46,47^. Based on assumptions about population size and demographic distribution of bats in this trial, an estimated gel uptake by 88.0% (CI: 76.9% - 94.0%) of the colony was measured after initial treatment of 20.5% of the colony. This corresponds to gel uptake in 3.29 bats (CI: 2.75 – 3.58) per initial bat treated which improves upon the transfer reported in Bakker et al. (2019) of 1.8-2.1^17^. This difference is likely related to the increase in volume of gel applied per bat from 0.45 mL to 1 mL and the higher proportion of remaining non-treated bats available for gel uptake. Given the near 100% uptake observed in adult females and juveniles in our study, gel transfer rates may be even higher given the finite number of untreated bats within the colony.

Importantly, adult male bats are disproportionately implicated in the dispersal of VBR^48^. While our field trial suggests that adult male bats may be less likely to consume orotopically applied vaccines vehicles, we still demonstrate predicted uptake by 48.4% (CI: 30.4%-69.6%) of non-treated adult male bats. It has also previously been documented that males may be less effective than females at spreading vampiricide within a colony^32^. However, this result differs from Bakker et al. (2019) who found a nonsignificant difference in the proportion of males that showed uptake of RB compared to females^17^. This previous finding could be associated with the high proportion of treatment (up to 50%) in the colonies reported, variation in the social dynamics within the colonies between the studies, or potential regional or seasonal influences. Vampire bat colonies are differentiated as either maternity colonies with resident and non-resident males or primarily-male bachelor roosts^49,50^. Social grooming interactions and thus topical vehicle uptake may vary between different sites with different structures and sex and age ratios^51^. Notably, 2.3-9 times the proportion of juveniles (31.5%) were captured in the roost in the present study compared to the three roosts studied in Bakker et al. (2019) (3.5%, 8.7% and 13.6%) during the first diurnal capture^17^.

While RFID detection remained stable between captures 1 and 2 in this study, the number detected declined by around 50% a week later. In addition to the multiple captures at this site, disturbance was caused by the euthanasia of 20 of the bats from our colony carried out by authorities of the Mexican Rabies Campaign as equine rabies cases were reported near the roost site. As previously reported^13,17^, vampire bats are prone to roost abandonment when disturbed, relocating to nearby manmade and natural roosts. For example, five months before the initiation of this biomarker study, we visited a large colony of over 100 bats within an abandoned facility 10 km away as a potential study site. However, before captures could be initiated, the colony relocated with only a few bats remaining at the site.

Epidemiological models indicate that vaccination significantly reduces duration and frequency of rabies outbreaks compared to culling even at low rates of vaccination^17^.

The estimated gel consumption rates by nearly 100% of adult females, and male and female juveniles found in this study will likely aid in preventing rabies virus introduction to naive colonies. As a lower proportion of adult males consumed the gel, targeting males as juveniles may result in higher rates of vaccination; however, this will require investigation of the development and duration of the immune response in vaccinated juvenile bats. Further field trials are needed to understand gel uptake dynamics within multiple bat colonies to maximize vaccine uptake by adult males. Additional models may also investigate the effect of differing rates of vaccination uptake between males and females. Given that vampire bat rabies has been known to spread among colonies in a slow wavelike fashion^1,52,53^, strategies of ring vaccination around areas of predicted spread may be effective at containing expansion of VBR^17^.

CMC gel may also work well as a topical vehicle for vaccine delivery to other bat species. In the United States, an estimated 40,000 individuals receive rabies prophylaxis each year after exposure to various bat species^10^. Furthermore, 36 of 40 (90%) fatal rabies cases acquired in the United States between 2000 and 2022 were of bat origin^11^. CMC could additionally be used to deliver a white nose syndrome (WNS) vaccine^54^ or to scale up deployment of a herpesvirus-vectored rabies vaccine^55^. As WNS-affected bats often inhabit colder regions in the United States and Canada, it is notable that CMC additionally maintains its physical properties at colder temperatures. However, vampire bats participate in allogrooming at higher rates than other bat species^56^, and field studies would be needed to determine gel transfer within colonies of other targeted bat species.

The identification of vampire bats within 50 km of the southern border of the United States^57^ and the expectation that their geographic range will continue to expand emphasizes the need for superior control methods^7^. While topical application of vaccines presents an appealing alternative to culling practices that may exacerbate rabies transmission, further studies are needed to understand the dimorphism of uptake rates in adult male and female bats observed in this study. Likewise, additional studies are needed to determine the impact of vaccination on vampire bat colony sizes, although it is proposed to include contraceptives as a method for population control while minimizing unintended ecological impacts of anticoagulant poisons^58^.

## Conclusion

This study details the characterization of a carboxymethyl cellulose (CMC) gel vehicle for orotopical rabies vaccine delivery to vampire bats. The new CMC formulation maintains optimal physical properties across a broad range of temperatures improving on a past topical vehicle. CMC gel is also able to sufficiently preserve our recombinant viral-vector rabies vaccine candidate. A field study in Jalisco, Mexico demonstrates the capacity for high rates of uptake of the CMC-based delivery formulation within a wild vampire bat colony at a low level of application. In summary, CMC gel provides an orotopical vehicle for vaccine delivery to bat species that socially groom and may be tested to treat other diseases and species beyond rabies in vampire bats.

## Data Availability

The data and code that support the findings of this study are openly available in github at https://github.com/zamanianlab/Bat_vaccine_gel-ms.

## Acknowledgements

We are very grateful for the support of team members from the Universidad de Guadalajara - Centro Universitario de la Costa Sur (CUCSUR) and Comité Estatal para el Fomento y Protección Pecuaria Jalisco (CEFPPJ) who aided in the field work. We would also like to thank the Centro Nacional de Servicios de Diagnóstico en Salud Animal (CENASA) in Tecamac, Mexico inviting us to work with their fluorescence microscope and assisting with visualizing the hair samples.

The authors gratefully acknowledge the use of facilities and instrumentation supported by NSF through the University of Wisconsin Materials Research Science and Engineering Center (DMR-1121288, 0079983, 0520057, 0832760, and 0425880) with special thanks to staff of the Soft Materials Characterization Lab for providing training on rheometer use and data analysis.

The project was funded by the NSF/BBSRC Ecology and Evolution of Infectious Diseases Program (DEB 2011069, BB/V003798/1) and the University of Wisconsin Madison Comparative Biomedical Sciences Student Travel Award (C.K.). D.G.S. was funded by a Wellcome Trust Senior Research Fellowship (217221/Z/19/Z). This research received support from the UK Medical Research Council through core funding of the MRC-University of Glasgow Centre for Virus Research (Virus Cross Species Transmission Programme: MC_UU_00034/3).

Any use of trade, firm, or product names is for descriptive purposes only and does not imply endorsement by the U.S. Government.

## Author Contributions

Conceptualization, C.K., E.M.C.C., D.P.W., D.G.S., J.E.O., M.Z., T.E.O.; data curation, C.K., T.M.G., M.Z.; formal analysis, C.K., T.M.G., funding acquisition, C.K., D.P.W., D.G.S., J.E.O., M.Z., T.E.O.; investigation C.K., E.M.C.C., T.M.G., M.M.R.M., D.L., L.P., D.P.W., D.G.S, T.E.O.; methodology, C.K., E.M.C.C., D.L., M.Z., T.E.O.; project administration, C.K., E.M.C.C., M.M.R.M., D.P.W., D.G.S., M.Z., T.E.O.; resources, M.M.R.M, J.E.O., M.Z., T.E.O.; software, C.K., T.M.G.; supervision, E.M.C.C., M.M.R.M., M.Z., T.E.O.; validation, C.K., E.M.C.C., M.Z., T.E.O.; visualization, C.K., T.M.G., D.L.; writing – original draft preparation, C.K.; writing – review & editing, C.K., E.M.C.C., T.M.G., M.M.R.M., D.L., L.P., D.P.W., D.G.S., J.E.O., M.Z., T.E.O.

