## Supplemental Information for "An optimized rabies vaccine vehicle for orotopical administration to wild vampire bats"

***This information product has been peer reviewed and approved for publication as a preprint by the U.S. Geological Survey.***

**An optimized rabies vaccine vehicle for orotopical administration to wild vampire bats**

**Authors**

Cole Knuese^1^, Elsa M. Cárdenas-Canales^1^, Travis McDevitt-Galles^2^, María Magdalena Ramírez Martínez^3^, Daniel Limonta^1^, Lisa E. Powers^2^, Daniel P. Walsh^4^, Daniel G. Streicker^5,6^, Jorge E. Osorio^1,7^, Mostafa Zamanian^1^, Tonie E. Rocke^2*^

**Affiliations**

^1^ School of Veterinary Medicine, University of Wisconsin, Madison, Wisconsin, USA.

^2^ US Geological Survey National Wildlife Health Center, Madison, Wisconsin, USA.

^3^ Departamento de Ciencias de la Salud y Ecología Humana, Centro Universitario de la Costa Sur, Universidad de Guadalajara, Autlán, Jalisco, México.

^4^ U.S. Geological Survey, Montana Cooperative Wildlife Research Unit, Wildlife Biology Program, University of Montana, Missoula, Montana, 59812, USA.

^5^ School of Biodiversity, One Health and Veterinary Medicine, University of Glasgow, Glasgow, UK.

^6^ MRC-University of Glasgow Centre for Virus Research, Glasgow, UK.

^7^ Global Health Institute, University of Wisconsin, Madison, Wisconsin, USA.

*

Any use of trade, firm, or product names is for descriptive purposes only and does not imply endorsement by the U.S. Government.

**Supplemental Information**

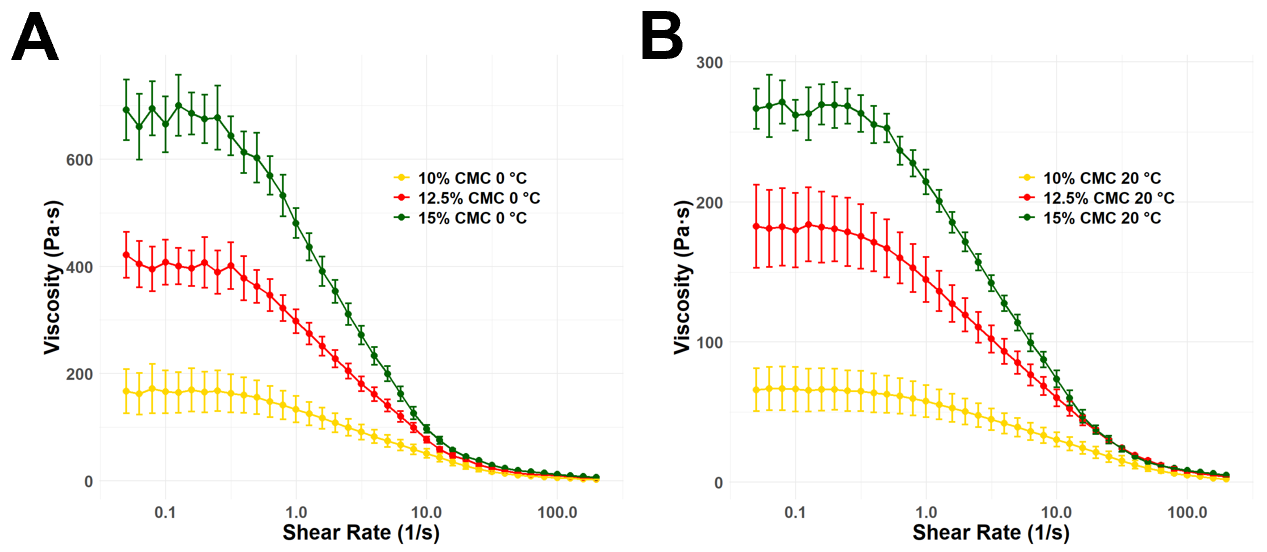

**Figure S1. Effect of carboxymethyl cellulose (CMC) concentration on viscosity at 0** °**C and 20** °**C**. Steady rate sweep test was performed with an ARES LS2 Advanced research grade rheometer (TA Instruments, New Castle, Delaware, USA) to measure viscosity of 10%, 12.5%, and 15% CMC at **A)** 0 °C and **B)** 20 °C. Error bars represent standard error of three experimental replicates.

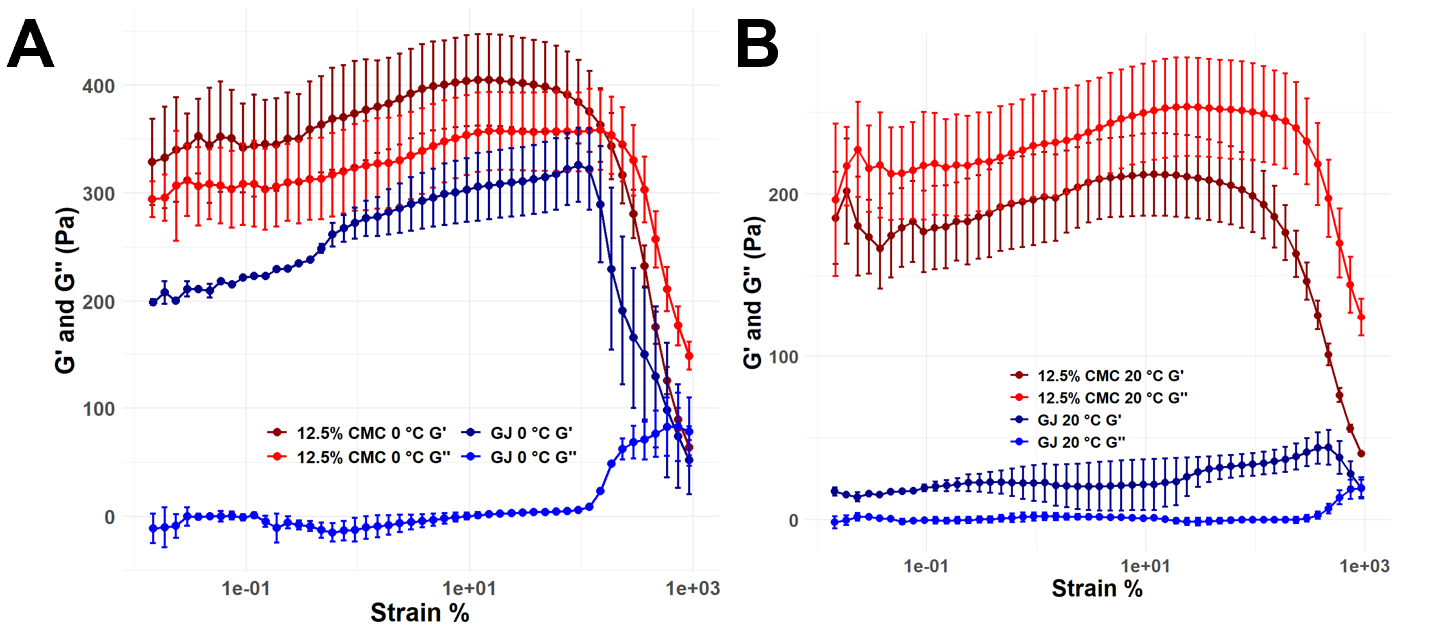

**Figure S2. Dynamic strain sweep test of 12.5% carboxymethyl cellulose (CMC) gel and glycerin jelly (GJ) at a frequency of 1 Hz at 0** °**C and 20** °**C.** Viscoelastic properties of CMC and GJ were measured by a dynamic strain sweep test at **A)** 0 °C and **B)** 20 °C. G’ represents the elastic modulus – energy stored during deformation, while G” represents the loss modulus – energy dissipated during deformation. All rheological measurements were recorded with an ARES LS2 Advanced research grade rheometer (TA Instruments, New Castle, Delaware, USA). Error bars represent standard error of three experimental replicates except for GJ at 0 °C where two replicates were captured.

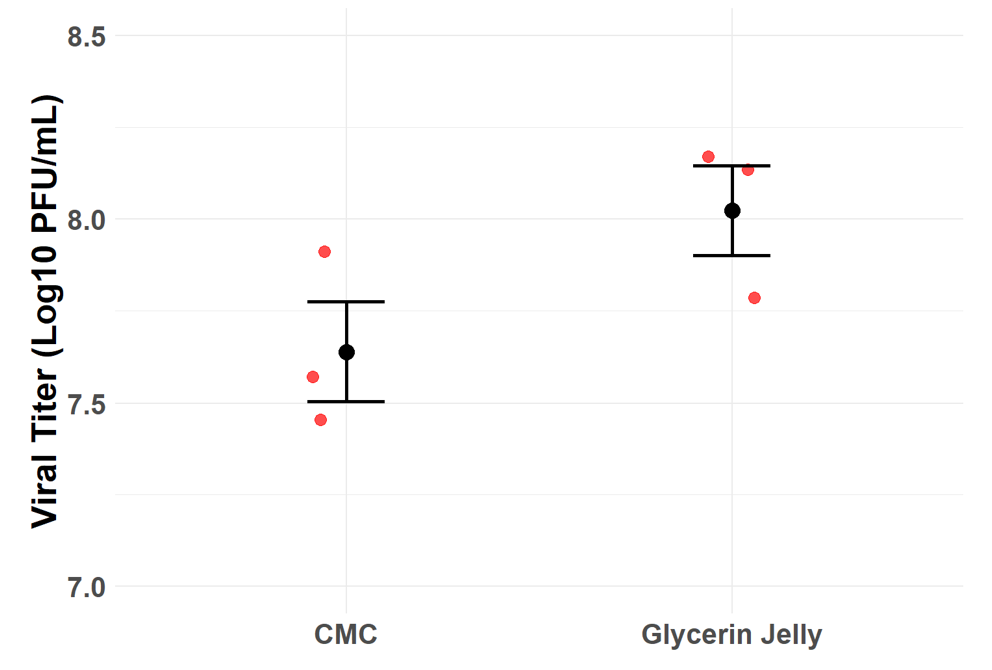

**Figure S3. Recovery of a recombinant raccoonpox vaccine expressing a mosaic lyssavirus glycoprotein (RCN-MoG) from carboxymethyl cellulose (CMC) gel and glycerin jelly (GJ) at T=0.** An initial titer of 10^8^ plaque forming units (PFU)/mL was mixed into CMC and GJ and frozen at -80 °C. The gels were thawed, dissolved, and titered by plaque assay on Vero cells (ATCC CCL-81). Red dots represent experimental replicates and error bars represent standard error.

**Real Time qPCR of stability samples with SYBR Green**

DNA was extracted from our recombinant raccoonpox vaccine expressing a mosaic lyssavirus glycoprotein (RCN-MoG) stored in CMC gel at T=0, and one month and three months at both 4 °C and 23 °C using Zymo Quick DNA Mini Prep kit (Zymo Research, Tustin, California, USA). Tables S1 and S2 were used for preparing the master mix and programming thermocycle conditions. The primer MoG-F targets the MoG protein, while RCN-R targets the raccoonpox vector.

**Table S1. SYBR green quantitative reverse transcription polymerase chain reaction (qPCR) master mix**

| **Reagent** | **Volume per sample (μL)** |
| --- | --- |
| Sso Advanced Universal SYBR Green Master Mix (Bio-Rad, Hercules, California, USA) | 10 |
| MoG-F primer (5’-GGAAGAGTAATATCTTCTTGGGAATC-3’) | 0.5 |
| RCN-R primer (5’CTATAACTATTTTTCCATTGTTTGCCATG-3’) | 0.5 |
| Nuclease free water | 7 |
| Sample DNA | 2 |
| **Total** | **20** |

**Table S2. SYBR green quantitative reverse transcription polymerase chain reaction (qPCR) thermocycle conditions**

| **Step** | **Temperature (**°**C)** | **Time** | **# of Cycles** |
| --- | --- | --- | --- |
| **Initial Denaturation** | 98 | 3 min | 1 |
| **Denaturation** | 98 | 30 s | 40 |
| **Annealing and Extension** | 60 | 20 s |  |

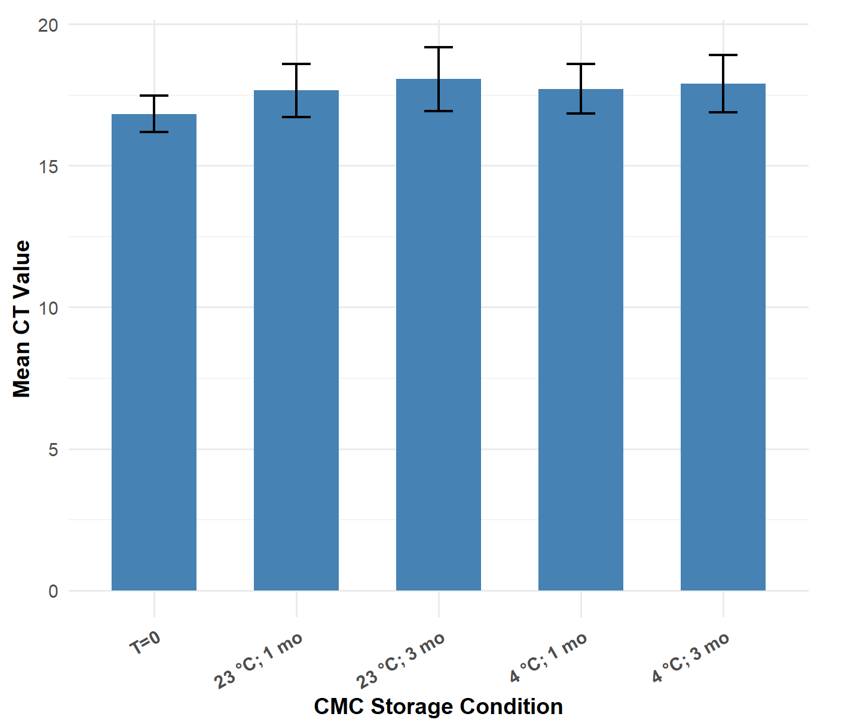

**Figure S4. Ct values from SYBR Green quantitative PCR measuring DNA stability of our recombinant raccoonpox vaccine expressing a mosaic lyssavirus glycoprotein (RCN-MoG) within carboxymethyl cellulose (CMC) gel in samples stored at 4** °**C and 23** °**C for one and three months.** Ct values were averaged across results of DNA extracted from all three experimental replicates of the RCN-MoG stability assay. Error bars represent the standard error. No statistical comparisons were significant by t-test.
